# Alpha-satellite RNA transcripts are repressed by centromere-nucleolus associations

**DOI:** 10.1101/2020.04.14.040766

**Authors:** Leah Bury, Brittania Moodie, Liliana S. McKay, Karen H. Miga, Iain M. Cheeseman

## Abstract

Centromeres play a fundamental role in chromosome segregation. Although originally thought to be silent chromosomal regions, centromeres are actively transcribed. However, the behavior and contributions of centromere-derived RNAs have remained unclear. Here, we used single-molecule fluorescence in-situ hybridization (smFISH) to detect alpha-satellite RNA transcripts in intact human cells. We find that alpha-satellite RNA smFISH foci fluctuate in their levels over the cell cycle and do not remain associated with centromeres, displaying localization consistent with other long non-coding RNAs. Our results demonstrate that alpha-satellite expression occurs through RNA Polymerase II-dependent transcription, but does not require centromere proteins and other cell division components. Instead, our work implicates centromere-nucleolar associations as the major factor regulating alpha-satellite expression. The fraction of nucleolar-localized centromeres inversely correlates with alpha-satellite transcripts levels, explaining variations in alpha-satellite RNA between cell lines. In addition, alpha-satellite transcript levels increase substantially when the nucleolus is disrupted. Together, our results are inconsistent with a direct, physical role for alpha-satellite transcripts in cell division processes, and instead support a role for ongoing transcription in promoting centromere chromatin dynamics. The control of alpha-satellite transcription by centromere-nucleolar contacts provides a mechanism to modulate centromere transcription and chromatin dynamics across diverse cell states and conditions.

## Introduction

Chromosome segregation requires the function of a macromolecular kinetochore structure to connect chromosomal DNA and spindle microtubule polymers. Kinetochores assemble at the centromere region of each chromosome. The position of centromeres is specified epigenetically-defined by the presence of the histone H3-variant, CENP-A, such that specific DNA sequences are neither necessary nor sufficient for centromere function. (McKinley and Cheeseman, 2016). However, despite the lack of strict sequence requirements, centromere regions are typically characterized by repetitive DNA sequences, such as the alpha-satellite repeats found at human centromeres. Understanding centromere function requires knowledge of the centromere-localized protein components as well as a clear understanding of the nature and dynamics of centromere chromatin. Although originally thought to be silent chromosome regions, centromeres are actively transcribed (Perea-Resa and Blower, 2018). Prior work has detected α-satellite transcription at centromere and pericentromere regions based on the localization of RNA polymerase II (Bergmann et al., 2012; Chan et al., 2012) and the production of centromere RNA transcripts (Chan et al., 2012; Saffery et al., 2003; Wong et al., 2007). Centromere transcription and the resulting RNA transcripts have been proposed to play diverse roles in kinetochore assembly and function (Biscotti et al., 2015; Blower, 2016; Fachinetti et al., 2013; Ferri et al., 2009; Grenfell et al., 2016; Ideue et al., 2014; McNulty et al., 2017; Quenet and Dalal, 2014; Rosic and Erhardt, 2016; Wong et al., 2007). However, due to limitations for analyses of centromere transcripts that average behaviors across populations of cells and based on varying results between different studies, the nature, behavior, and contributions of centromere-derived RNAs remain incompletely understood.

Here, we used single-molecule fluorescence in-situ hybridization (smFISH) to detect alpha-satellite RNA transcripts in individual, intact human cells. Our results define the parameters for the expression and localization of centromere and pericentromere-derived transcripts across a range of conditions. We find that the predominant factor controlling alpha-satellite transcription is the presence of centromere-nucleolar contacts, providing a mechanism to modulate centromere transcription and chromatin dynamics across diverse cell states and conditions. Together, our results support a role for ongoing transcription in promoting the dynamics of centromeric chromatin (Swartz et al., 2019).

## Results and Discussion

### Quantitative detection of alpha-satellite RNAs by smFISH

Prior work has analyzed centromere RNA transcripts primarily using population-based assays, such as qPCR and RNA-seq, or has detected centromere RNAs in spreads of mitotic chromosomes. To visualize alpha-satellite RNA transcripts in individual intact human cells, we utilized single-molecule fluorescence in-situ hybridization (smFISH), a strategy that has been used to detect mRNAs and cellular long non-coding RNAs (lncRNAs) (Raj et al., 2008). The high sensitivity of smFISH allows for the accurate characterization of number and spatial distribution of RNA transcripts.

Alpha-satellite DNA is degenerate such that it can vary substantially between different chromosomes with the presence of higher order repeats of alpha-satellite variants (Waye and Willard, 1987) (Willard and Waye, 1987b). Thus, we first designed targeted probe sets to detect RNAs derived from centromere regions across multiple chromosomes: 1) Sequences complementary to a pan-chromosomal consensus alpha-satellite sequence (labeled as “ASAT”), 2) sequences that target Supra-Chromosomal family 1 (SF1) higher-order arrays, present on chromosomes 1, 3, 5, 6, 7, 10, 12, 16, and 19 (labeled as “SF1”) (Alexandrov et al., 2001; Uralsky et al., 2019), and 3) sequences that are enriched for transcripts from the Supra-Chromosomal family 3 higher-order arrays present on chromosomes 1, 11, 17 and X (labeled as “SF3”), with an increased number of targets on chromosome 17 (D17Z1) (Willard and Waye, 1987a). Second, we designed probes that detect sequences enriched on specific chromosomes including the X chromosome (DXZ1, labeled “X”; (Miga et al., 2014; Willard et al., 1983)) and Chromosome 7 (D7Z2, labeled as “7.2”; (Waye et al., 1987)). For complete sequence information and an analysis of sequence matches to different chromosomes, see the Experimental Procedures and Supplemental Table 2. Alpha-satellite DNA can span megabases of DNA on a chromosome, whereas the active centromere region is predicted to be as small as 100 kb in many cases (McKinley and Cheeseman, 2016). Thus, our smFISH probes will detect RNA transcripts from both the active centromere region and the flanking pericentric alpha-satellite DNA.

In asynchronously cycling HeLa cells, we detected clear foci using smFISH probe sets for ASAT, SF1, and SF3 (Fig. 1A). In contrast, we did not detect foci using oligos designed to recognize transcripts derived from the centromere regions of chromosome 7 or the X chromosome (Fig. 1B). As the absence of signal could reflect a variety of technical features of probe design or a detection limit for the expression level or length of these sequences, we chose not to pursue these probes further. For the probes that displayed clear foci, we sought to ensure that this signal was not due to non-specific hybridization of the RNA probes to genomic DNA. To test this, we treated cells with RNase A prior to hybridization. The RNA FISH signal was diminished substantially after RNase A treatment (Fig. 1A, C), confirming the ribonucleic source of the observed signal. To quantify the number of distinct RNA-FISH foci, we used CellProfiler (Carpenter et al., 2006) to measure the number of RNA foci per nucleus systematically using z-projections of the acquired images. The number of smFISH foci varied between individual cells, but averaged approximately 4 foci/cell for the ASAT, SF1, and SF3 probe sets in HeLa cells (Fig. 1C).

**Figure 1.**
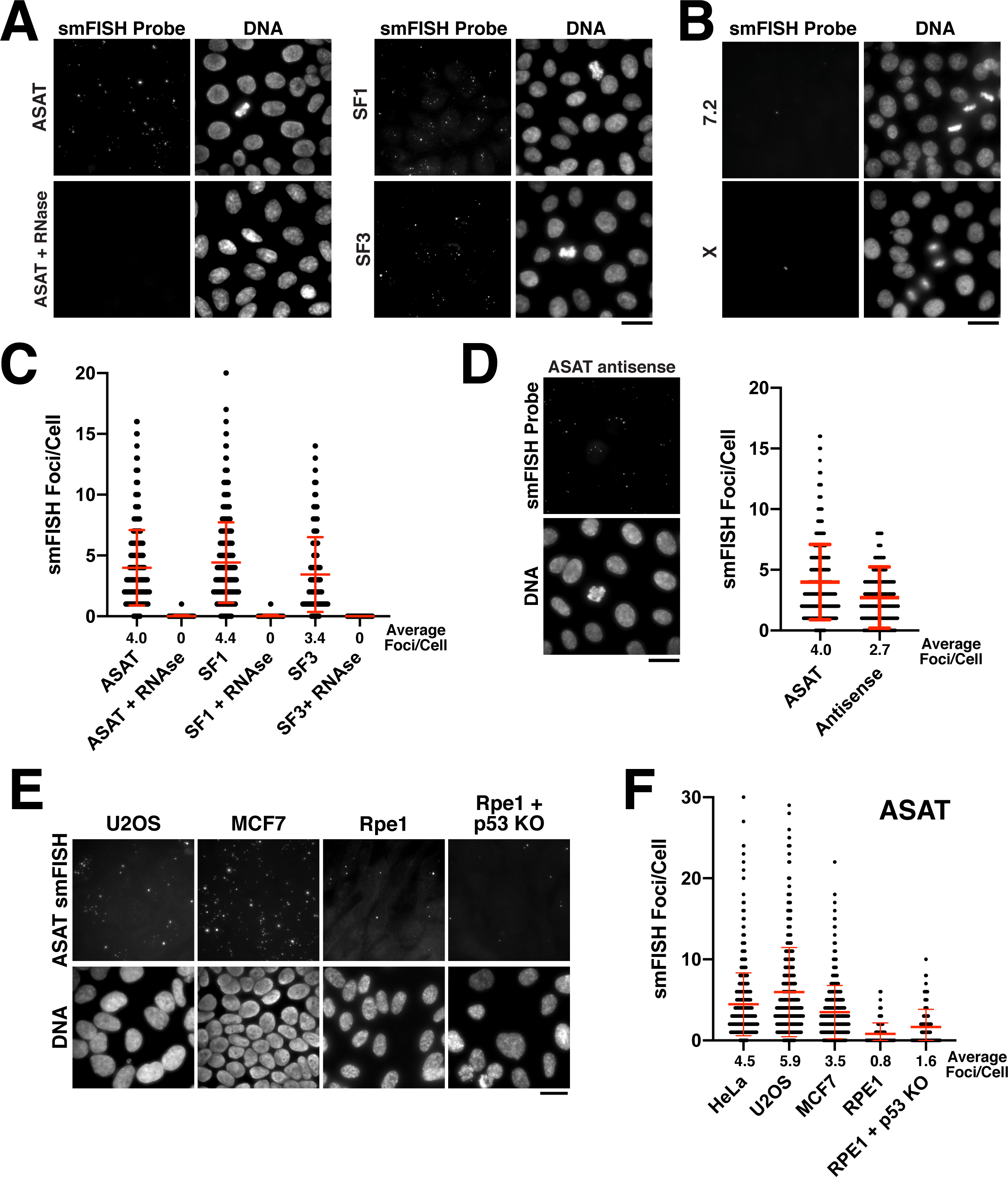
Quantitative detection of centromere RNAs using smFISH. A) Detection of alpha-satellite RNA transcripts by smFISH in asynchronous HeLa cells. Designed probes detected RNAs derived from centromeres across multiple chromosomes (ASAT) or specific chromosomes (3 and 17). Treatment of cells with RNase A prior to hybridization diminished RNA-smFISH signals. B) RNA transcripts are not detected using probes designed against centromere sequences from chromosome 7 and X. C) Quantification of smFISH foci in the presence or absence of RNase A treatment indicates that the signal observed is due to a ribonucleic source. Points represent the number of foci in a single nucleus of a cell. Error bars represent the mean and standard deviation of at least 100 cells. D) Detection of anti-sense alpha-satellite transcripts in HeLa cells for the ASAT smFISH probe sequences. Error bars represent the mean and standard deviation of at least 100 cells. E) Transcription of centromeric and pericentromeric satellite DNA varies across cell line, with RPE-1 cells displaying overall lower levels of centromere smFISH foci. F) Quantification indicating the variation of smFISH foci across selected cell lines. Quantification of an RPE1 cell line lacking the tumor suppressor p53 reveals that factors besides the cellular transformation status likely affect the cellular levels of centromere RNA transcripts. Error bars represent the mean and standard deviation of at least 100 cells. Scale bars, 25 µm

Transcription of non-coding RNAs often occurs from both strands of DNA at a given locus. We therefore tested whether we could detect antisense (relative to the “sense” probes used above) alpha-satellite transcripts in Hela cells using smFISH. Indeed, for the ASAT probe sequences, we were able to visualize ∼3 foci/cell using antisense smFISH probes, similar to numbers using the sense probe set (4 foci/cell) (Fig. 1D). Antisense transcription at the centromere has also been previously reported across a variety of species (Carone et al., 2009; Choi et al., 2011; Chueh et al., 2009; Ideue et al., 2014; Koo et al., 2016; Li et al., 2008; May et al., 2005).

The level of transcription of centromeric and pericentromeric satellite DNA has been proposed to vary between developmental stages and tissue types (Maison et al., 2010; Pezer and Ugarkovic, 2008). In addition, changes in centromere and pericentromere transcription have been observed in cancers (Ting et al., 2011). Therefore, we next sought to analyze differences in smFISH foci across different cell lines using the ASAT and SF1 probe sets. We selected the chromosomally-unstable osteosarcoma cell line U2OS, the breast cancer cell line MCF7, and the immortalized, but non-transformed hTERT-RPE-1 cell line. We found that the levels of alpha-satellite transcripts varied modestly across cell lines (Fig. 1E,F; Supplemental Fig. 1A,B), with RPE-1 cells displaying overall lower levels of centromere smFISH foci. To test whether the transformation status of the cell line correlated with the level of smFISH foci, we eliminated the tumor suppressor p53 in RPE-1 cells using our established inducible knockout strategy (McKinley and Cheeseman, 2017). Eliminating p53 did not substantially alter the levels of alpha-satellite smFISH foci in Rpe1 cells (Fig. 1E,F) indicating that other factors likely contribute to the cellular levels of centromere RNA transcripts. Together, this strategy provides the ability to quantitatively detect centromere and pericentromere-derived alpha-satellite RNA transcripts using smFISH probes against alpha-satellite sequences from different chromosomes and demonstrates that human cell lines display varying levels of alpha-satellite transcripts.

### Analysis of alpha-satellite transcript localization and cell cycle control

We next sought to assess the localization of alpha-satellite RNA transcripts within a cell. Prior work suggested that non-coding centromere transcripts are produced in *cis* and remain associated with the centromere from which they are derived, including through associations with centromere proteins (McNulty et al., 2017). Other studies support the action of centromere-derived RNAs in *trans* (Blower, 2016), but again acting at centromeres. To investigate the distribution of the centromere transcripts, we performed combined immunofluorescence and smFISH to visualize alpha-satellite transcripts relative centromeres and microtubules. In interphase cells, smFISH foci localized within the nucleus (Fig. 2A). Thus, unlike spliced mRNAs, alpha-satellite-derived RNAs are not exported to the cytoplasm. Although we detected co-localization of alpha-satellite RNAs with a subset of centromeres in HeLa cells, the majority of smFISH foci did not co-localize with centromeres (Fig. 2A). As cells entered mitosis, we observed the disassociation of smFISH foci from chromatin (Fig. 2B). During all stages of mitosis, alpha-satellite RNA transcripts appeared broadly distributed within the cytoplasm. Finally, as the cells exited mitosis into G1, the smFISH foci remained distinct from the chromosomal DNA and were thus excluded from the nucleus when the nuclear envelope reformed (Fig. 2C). Similar patterns of cell-cycle dependent localization changes with mitotic exclusion from chromatin have been reported for other cellular long non-coding RNAs (Cabili et al., 2015; Clemson et al., 1996). Thus, alpha-satellite-derived non-coding RNAs visualized by smFISH display nuclear localization, but are not tightly associated with the centromere regions from which they are derived.

**Figure 2.**
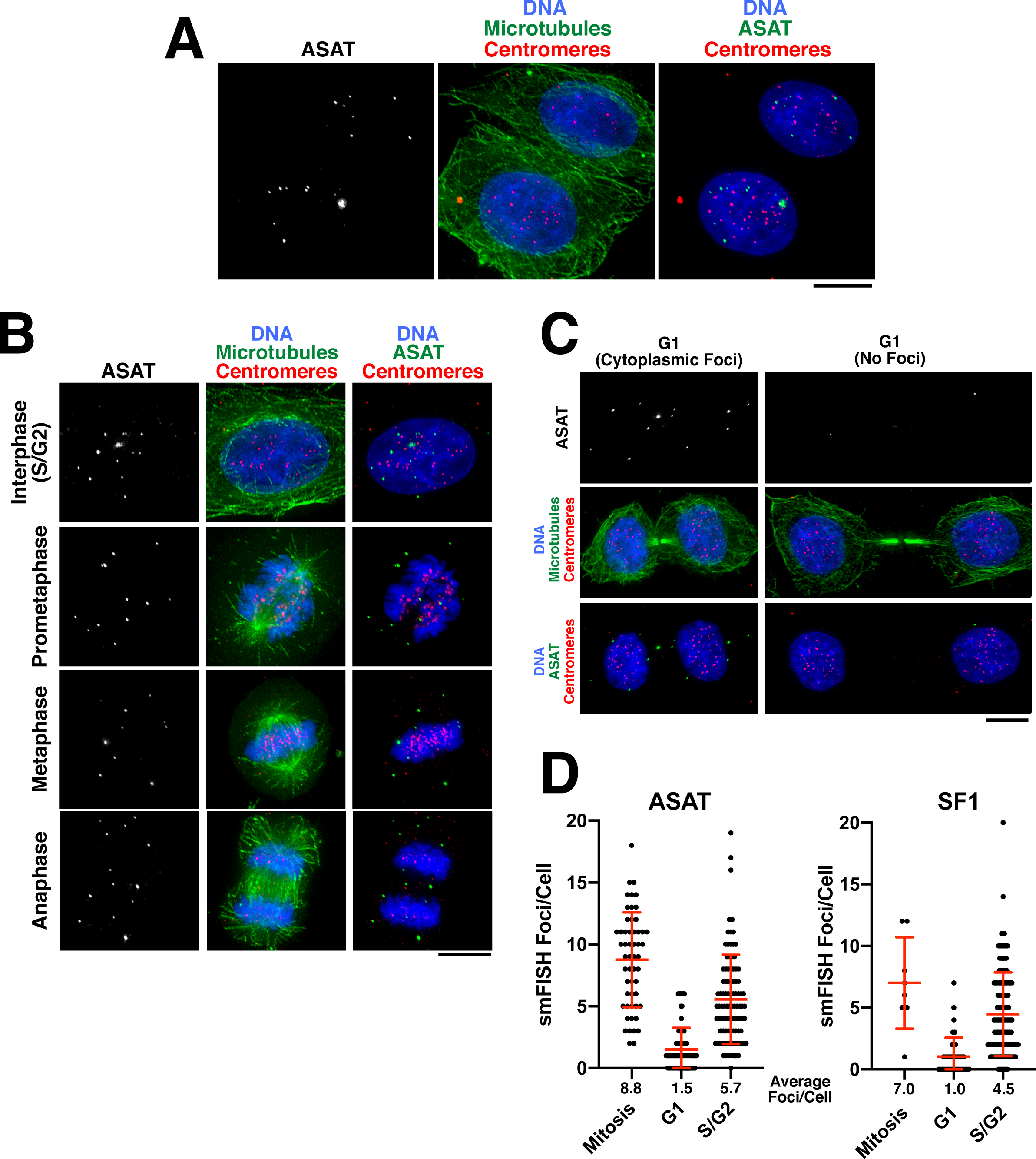
Analysis of centromere RNA foci across the cell cycle. A) Immunofluorescence images (using anti-tubulin antibodies in green and anti-centromere antibodies (ACA) in red) showing alpha-satellite derived transcripts (smFISH; ASAT probe sets) localized to the nucleus during interphase in HeLa cells. The majority of detected transcripts do not co-localize with centromeres. B) Immunofluorescence of HeLa cells (as in A) throughout the cell cycle reveals the disassociation of smFISH foci from chromatin in mitosis. C) Immunofluorescence-smFISH analysis indicates that progression of cells into G1 (defined by cells with a mid-body) results in the nuclear exclusion of smFISH foci. Left: Foci are located in the cytoplasm after the nuclear envelope reforms. Right: Foci are absent, possibly reflecting the degradation of cytoplasmic RNA. D) Quantification of smFISH foci throughout the cell cycle (for either ASAT or SF1 probe sets) reveals that transcripts levels are high in S/G2 and mitotic cells, but reduced as cells exit mitosis into G1. Error bars represent the mean and standard deviation of at least 8 cells. Scale bars, 10 µm.

We next analyzed the temporal changes in alpha-satellite transcript numbers during the cell cycle. In contrast to other genomic loci, RNA Polymerase II is present at human and murine centromeres during mitosis (Chan and Wong, 2012; Perea-Resa et al., 2020). In addition, centromere transcription during G1 has been proposed to play a role in CENP-A loading (Quenet and Dalal, 2014). Recent work measuring the levels of satellite transcripts originating from specific centromeres in human cells suggested the presence of stable RNA levels during the entire cell cycle (McNulty et al., 2017). smFISH provides the capacity to measure the levels of cenRNA transcripts in individual cells over the course of the cell cycle. We utilized combined immunofluorescence-smFISH to simultaneously label alpha-satellite RNA transcripts and microtubules, allowing us to distinguish between G1 cells (due to the presence of a midbody), an S/G2 interphase population, and mitotic cells. In contrast to previous observations, our analysis revealed that the transcripts detected by our smFISH method increased in S/G2 and remained stable throughout mitosis (Fig. 2D). We note that a G2/M peak of transcript levels has been reported for murine Minor Satellite transcripts (Ferri et al., 2009). However, as cells exited mitosis into G1, transcripts detected by smFISH were reduced (Fig. 2D). We speculate that this may result from the nuclear exclusion of the existing alpha-satellite transcripts, which would make this more susceptible to degradation by cytoplasmic RNAses. Thus, alpha-satellite transcript levels fluctuate over the cell cycle with G1 as a period of low transcript numbers, either indicating reduced transcription during this cell cycle stage or the increased elimination of alpha-satellite-derived RNA transcripts.

### Alpha-satellite RNAs are products of Pol II-mediated transcription

Previous studies have suggested that centromeres are actively transcribed by RNA polymerase II. RNA polymerase II localizes to centromeres in *S. pombe, Drosophila melanogaster*, and human cells, including at centromeric chromatin on human artificial chromosomes (HACs) and at neocentromeres (Bergmann et al., 2011; Catania et al., 2015; Chan and Wong, 2012; Chueh et al., 2009; Ferri et al., 2009; Li et al., 2008; Ohkuni and Kitagawa, 2011; Perea-Resa et al., 2020; Quenet and Dalal, 2014; Rosic et al., 2014; Wong et al., 2007). However, in addition to RNA polymerase II, RNA polymerase III has been implicated in the transcription of pericentromeric DNA (Scott et al., 2006; Scott et al., 2007). To determine the polymerases that are responsible for generating the alpha-satellite transcripts detected by our smFISH assay, we treated Hela cells with small molecule inhibitors against all three RNA polymerases. We found a significant reduction in alpha-satellite smFISH foci following inhibition of RNA Polymerase II activity using the small molecule THZ1 (Fig. 3A,B), which targets the RNA Pol II activator Cdk7 (Kwiatkowski et al., 2014). In contrast, we did not detect a reduction in smFISH foci following treatment with inhibitors against RNA polymerase I (small molecule inhibitor BHM-21; (Colis et al., 2014)) or RNA polymerase III (ML-60218; (Wu et al., 2003)) (Fig. 3A,B; Supplemental Fig. 2A,B). Instead, as discussed below, we found dramatically increased alpha-satellite smFISH foci following RNA polymerase I inhibition. This suggests that the alpha-satellite RNA transcripts detected by smFISH are products of RNA Pol II-mediated transcription.

**Figure 3.**
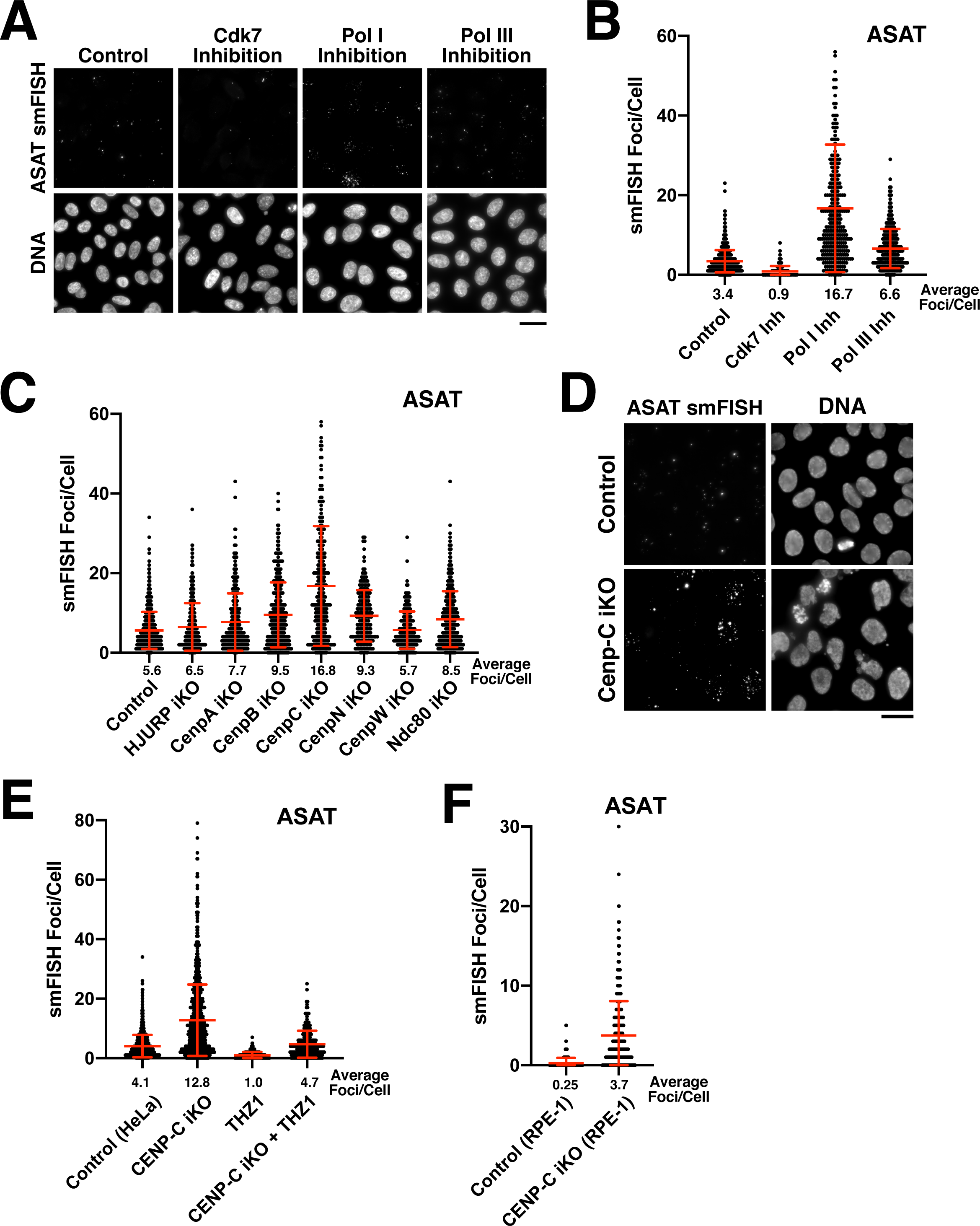
Functional analysis of the requirements for centromere RNA production. A) Treatment of HeLa cells with small molecule inhibitors reveals that alpha-satellite transcripts are mediated by RNA polymerase II. Cells were treated with the RNA Polymerase I inhibitor BMH-21, the RNA Polymerase III inhibitor ML-60218 or the Cdk7 inhibitor THZ1, which inhibits RNA Polymerase II initiation. Transcripts were identified using the ASAT smFISH probe sets. B) Quantification of smFISH foci after treatment of HeLa cells with small molecule inhibitors against Cdk7, RNA Pol I, and RNA Pol III. smFISH foci were significantly reduced after inhibition of RNA Pol II activator, Cdk7, but increased by RNA Pol I inhibition. Error bars represent the mean and standard deviation of at least 240 cells. C) Quantification of smFISH foci (ASAT probe set) after elimination of selected centromere and kinetochore components reveals that centromere components are not required for the specific recruitment of RNA Polymerase II to centromeres. Notably, CENP-C inducible knockout cells displayed a substantial increase in smFISH foci. Error bars represent the mean and standard deviation of at least 240 cells. D) Representative images of the substantial increase in smFISH foci after elimination of the centromere component CENP-C. E) The increase in alpha-satellite transcripts in cells depleted for CENP-C depends on RNA Polymerase II as THZ1 treatment resulted in a substantial reduction in smFISH foci in both control cells and CENP-C inducible knockout cells. F) Quantification of smFISH foci in CENP-C inducible knockout RPE-1 cells reveals that the increase in alpha-satellite transcripts following CENP-C knockout is not specific to HeLa cells. Error bars represent the mean and standard deviation of at least 170 cells. Scale bars, 25 µm.

### CENP-C acts to repress alpha-satellite RNA levels

We next sought to determine the requirements for the production of alpha-satellite transcripts. Centromere DNA functions as a platform for assembly of the kinetochore structure (McKinley and Cheeseman, 2016), an integrated scaffold of protein interactions that mediates the connection between the DNA and microtubules of the mitotic spindle. One possibility to explain the observed transcription of centromere regions, including at neocentromere loci lacking alpha-satellite sequences, is that centromere and kinetochore components could act to recruit the RNA Polymerase machinery. To test this, we selectively eliminated diverse centromere and kinetochore components using a panel of CRISPR inducible knockout cell lines expressing dox-inducible Cas9 and guide RNAs (McKinley and Cheeseman, 2017). For centromere proteins, we targeted the centromere-specific H3 variant CENP-A, the CENP-A chaperone HJURP (to block new CENP-A incorporation), the centromere alpha-satellite DNA binding protein CENP-B, and the constitutive centromere components CENP-C, CENP-N, and CENP–W. We also eliminated the outer kinetochore microtubule-binding protein Ndc80. Eliminating these factors did not prevent the presence of alpha-satellite RNA smFISH foci (Fig. 3C). In contrast, the number of foci/cell increased in many of these inducible knockout cell lines, from moderate increases in most knockout cell lines to a substantial increase in CENP-C inducible knockout cells (Fig. 3C). This suggests that centromere components are not required for the specific recruitment of RNA Polymerase II to centromere regions, although active centromeres may act to retain RNA Polymerase II during mitosis due to the persistence of sister chromatid cohesion (Perea-Resa et al., 2020).

We also tested the contribution of non-centromere localized cell division components to alpha-satellite transcription. Because of its DNA-based nature, the centromere is subject to cell cycle-specific challenges that include chromatin condensation, cohesion, and DNA replication. We thus sought to assess whether disruption of any of these complexes would influence alpha-satellite RNA transcript levels. To do this, we targeted proteins involved in centromere regulation (Sgo1 and BubR1), DNA replication (Mcm6, Gins1, Orc1, and Cdt1), sister chromatid cohesion (ESCO2, Scc1), chromosome condensation (Smc2, CAPG, CAPG2, TOP2A), and nucleosome remodeling (SSRP1). Strikingly, despite the diverse roles of these proteins in different aspects of centromere function, none of these inducible knockouts resulted in reduced levels of ASAT alpha-satellite transcripts as detected by smFISH analysis (Supplemental Fig. 2C). Instead, in many cases we detected an increase in alpha-satellite smFISH foci in the inducible knockout cells.

Of proteins that we tested, eliminating CENP-C had a particularly substantial effect on the number of smFISH foci. To confirm this behavior following the loss of CENP-C, we repeated these experiments for both the ASAT and SF1 smFISH probes (Fig. 3D,E; Supplemental Fig. 2D). In both cases, we observed a strong increase in smFISH foci. To test whether this behavior was specific to HeLa cells, we analyzed the CENP-C inducible knockout in RPE-1 cells. Although there are fewer ASAT smFISH foci in the parental RPE-1 cells, eliminating CENP-C resulted in a strong increase in the number of ASAT smFISH foci (Fig. 3F). This increase in alpha-satellite transcripts in cells depleted for CENP-C depends on RNA Polymerase II, as THZ1 treatment resulted in a substantial reduction in smFISH foci for the ASAT and SF1 probe sets in both control cells and CENP-C inducible knockout cells (Fig. 3E; Supplemental Fig. 2D). As eliminating CENP-C potently disrupts the localization of all centromere proteins (McKinley et al., 2015), this suggests that centromere and kinetochore formation could act as a physical block to restrict the passage of RNA polymerase through the centromere, downregulating alpha-satellite transcript levels. Alternatively, kinetochore proteins could act to create a repressive environment for transcription (see below).

Overall, our results indicate alpha-satellite transcription does not require the presence of specific DNA binding proteins or DNA structures, and instead that multiple factors act to restrict transcription at centromeres.

### Nucleolar associations act to repress alpha-satellite transcription

In the functional analysis described above, we were surprised that most perturbations resulted in increased centromere smFISH RNA foci instead of a loss of signal. The largest increases were observed for the depletion of CENP-C (Fig. 3C,E) and the inhibition of RNA Polymerase I (Fig. 3B). RNA Polymerase I transcribes rDNA, but also has an important role in assembling the nucleolus, which creates a repressive transcriptional environment. Given reported connections between the centromere and nucleolus in prior work (Ochs and Press, 1992; Padeken et al., 2013; Wong et al., 2007), we hypothesized that alpha-satellite transcription occurs at a basal level, but that this transcription is repressed by associations between the centromere and the nucleolus. To test this model, we first visualized centromeres and nucleoli in human cells. In HeLa and RPE-1 cells, a subset of centromeres overlap with the nucleolus, as marked with antibodies against Ki-67 (Fig. 4A) or Fibrillarin (Supplemental Fig. 3A). However, other centromeres are present outside of the nucleoli within the rest of the nucleus. This contrasts with work in Drosophila cells, where centromeres from all four chromosomes are found in close proximity surrounding the nucleolus (Padeken et al., 2013). Importantly, we observed an inverse relationship between the fraction nucleoli-localized centromeres and the numbers of alpha-satellite smFISH foci. First, we observed an increased fraction of nucleoli-localized centromeres in RPE-1 cells compared to HeLa cells (Fig. 4A,B), correlating with the reduced numbers of alpha-satellite smFISH foci in RPE-1 cells (Fig. 1F). Similarly, we found that CENP-C inducible knockout cells displayed a reduced fraction of nucleoli-localized centromeres (Fig. 4C,D), again correlating with the increased alpha-satellite smFISH foci in these cells (Fig. 3E). When the different conditions affecting the nucleolus are compared, there is a clear inverse relationship between nucleolar-localized centromeres and the number of ASAT smFISH foci per cell (Fig. 4E).

**Figure 4.**
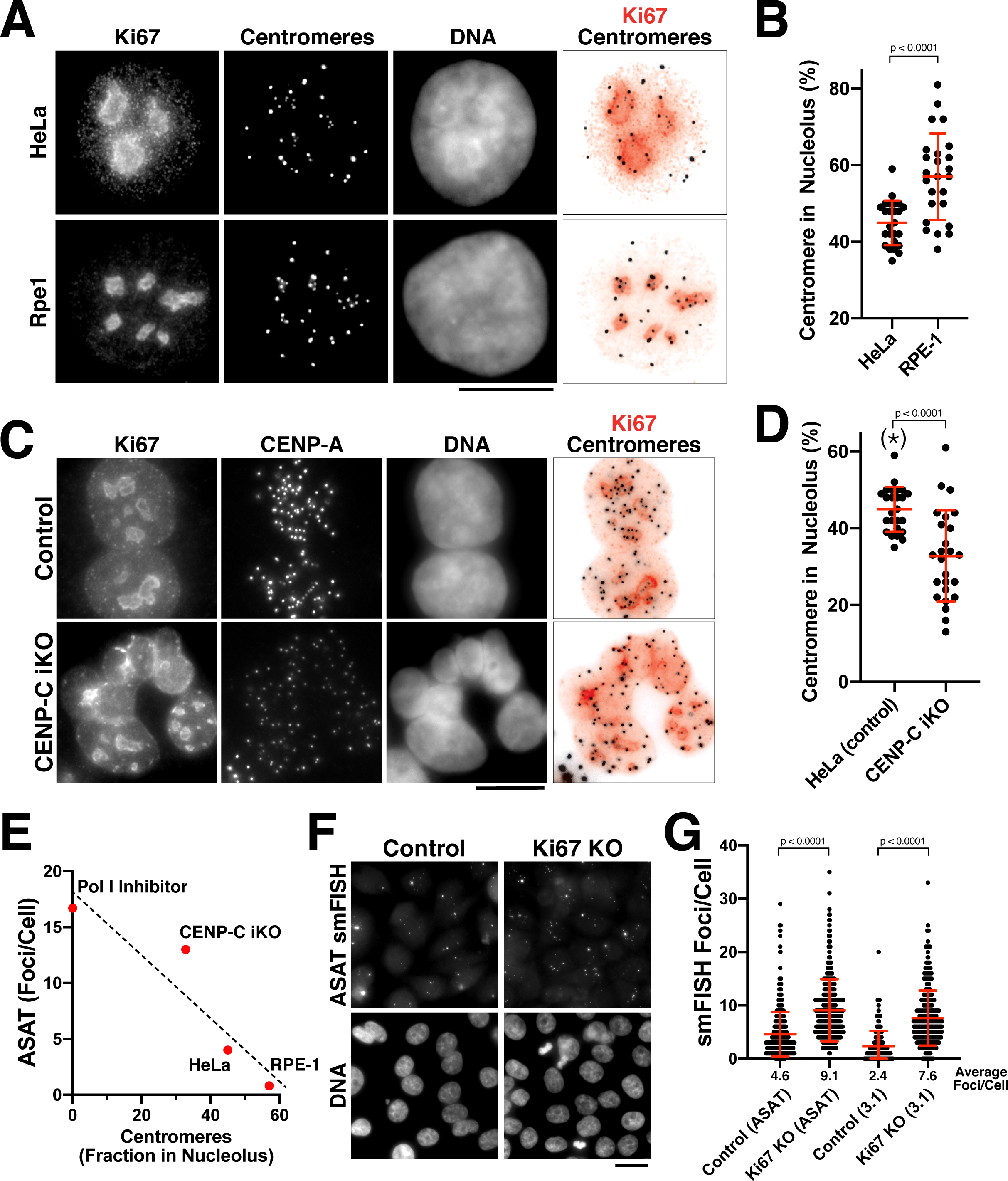
The nucleolus represses centromere RNA production. A) Immunofluorescence of HeLa (Top) and RPE1 (Bottom) cells showing the colocalization of centromeres with the nucleolus, as marked with antibodies against Ki-67 and anti-centromere antibodies (ACA). Scale bars, 10 µm. B) Quantification reveals RPE1 cells have a greater averge percentage of centromere overlap with nucleoli (57%) compared to HeLa cells (44.6%). Error bars represent the mean and standard deviation of 25 cells. C) Immunofluorescence of HeLa control (top) and HeLa CENP-C iKO (bottom) cells showing the colocalization of centromeres with the nucleolus, as marked with antibodies against Ki-67 and CENP-A. Scale bar, 10 µm. D) Quantification reveals that depletion of CENP-C results in a reduced fraction of nucleoli-localized centromeres (32.8%) compared to control cells (44.6%). The asterisk indicates that the data from control cells is repeated from (B). Error bars represent the mean and standard deviation of 25 cells. E) Graph showing the relationship between the number of ASAT smFISH foci and the fraction of nucleolar-localized centromeres in the indicated conditions (summarized from data in Figures 3 and 4). RNA Polymerase I inhibition should eliminate nucleolar function, and so is listed as “0” for nucleolar centromeres. Dashed line shows a linear fit trendline. F) smFISH assay reveals an increase of alpha-satellite transcripts in Ki67 knockout cells (right) when compared to control (left). Scale bar, 25 µm. G) Quantification reveals a 2-3 fold increase in alpha-satellite transcript levels for both the ASAT and SF1 smFISH probes in Ki67 knockout cells. Based on the increased alpha-satellite transcript levels when Ki-67 are inhibited, a properly functioning nucleolus is required to limit centromere and pericentromere transcription. Error bars represent the mean and standard deviation of at least 100 cells. Where indicated, p values indicate unpaired t-tests.

To assess the functional relationship between the nucleolus and alpha-satellite transcription, we generated inducible knockout cell lines for the nucleolar components Fibrillarin and Ki-67 using our established inducible Cas9 knockout system (McKinley and Cheeseman, 2017). Induction of these knockouts resulted in increased levels of alpha-satellite transcription, particularly for Ki-67 (Supplemental Fig. 3B). Although Ki-67 plays important roles in nucleogenesis and mitotic chromosome structure (Booth et al., 2014; Cuylen et al., 2016; Sobecki et al., 2016), deletion of Ki-67 is not lethal (Sobecki et al., 2016). Therefore, we additionally generated a stable Ki67 knockout cell line in HeLa cells (Supplemental Fig. 3C). Ki-67 knockout cells proliferated normally, but displayed a 2-3 fold increase in alpha-satellite transcript levels for both the ASAT and SF1 smFISH probes (Fig. 4F,G). However, despite these persistently increased alpha-satellite transcript levels, we did not detect notable consequences to centromere protein levels (based on the localization of CENP-A; Supplemental Fig. 3C) or chromosome mis-segregation (not shown). Thus, based on the increased alpha-satellite transcript levels when RNA Polymerase I or Ki-67 are inhibited, a properly functioning nucleolus is required to limit centromere and pericentromere transcription.

Together, this work defines the parameters for the production of alpha-satellite RNA transcripts and demonstrates that centromere-nucleolar connections act to restrict alpha-satellite transcription. The nature of the behavior that we observed for alpha-satellite smFISH foci, including the lack of localization to centromeres or mitotic structures, is inconsistent with a direct, physical role for these transcripts in cell division processes. Instead, this suggests a role for the transcription process itself, such as a role for ongoing transcription to promote the dynamics and rejuvenation of centromeric chromatin, as proposed by our recent work in non-dividing cells (Swartz et al., 2019). Together, we propose that basal centromere transcription acts to promote the turnover of DNA-bound proteins, providing a mechanism to ensure the rejuvenation of CENP-A chromatin. Importantly, changes in nuclear and nucleolar organization and in centromere-nucleolar associations across cell types and between cell states has the potential to create consequential changes to centromere transcription and centromere protein dynamics.

## Acknowledgements

We thank the members of the Cheeseman lab, Oceane Marescal, and Gayathri Muthukumar for their support and input. This work was supported by grants from The Harold G & Leila Y. Mathers Charitable Foundation, the NIH/National Institute of General Medical Sciences (GM108718 and R35GM126930) to IMC, and an American Cancer Society post-doctoral fellowship to LB. The authors declare that they have no conflict of interest.

## Author Contributions

Conceptualization – LB, KM, IMC; Methodology – LB, BM, LSM, KM, IMC; Validation – LB, BM LSM; Investigation – LB, BM, LSM, IC; Writing - Original Draft Preparation – IMC; Writing – Review & Editing – LB, BM, LSM, KM, IMC; Visualization – SZS, LSM, KCS, IMC; Supervision: IMC; Funding Acquisition: IMC.

## Methods

### Cell Culture

All cells were grown in Dulbecco’s Modified Eagle’s Medium (DMEM) supplemented with 10% Fetal Bovine Serum, 100 units/mL penicillin, 100 units/mL streptomycin, and 2 mM L-glutamine (Complete Media) at 37°C with 5% CO_2_. For experiments using inducible knockout cell lines, cells were seeded onto uncoated glass coverslips and doxycycline (DOX, Sigma) was added at 1 mg/L for 48 hours. Cells were fixed and stained at 4 or 5 days following DOX addition. For inhibitor experiments, RNA Polymerase inhibitors were added to cells for 24 h at the following concentrations: BMH-21 (RNAPI inhibitor; Millipore) at 1 µM; ML-60218 (RNAP III inhibitor; Fisher) at 20 µM; THZ1 (Cdk7 Inhibitor; Fisher Scientific) at 1 µM.

Inducible knockouts for nucleolar components in HeLa cells were created as described previously (McKinley and Cheeseman, 2017). Briefly, sgRNA sequences were cloned into pLenti-sgRNA (Wang et al., 2015), and used to generate lentiviruses for stable infection in cells harboring inducible Cas9 (HeLa cells – cTT20; RPE-1 cell - cTT33). Cells were then selected with puromycin as described previously (McKinley and Cheeseman, 2017). Additional cell cycle and chromosome inducible knockouts were from (McKinley and Cheeseman, 2017). For the Ki67 stable knockout cell line, the HeLa cell inducible knockout version was induced with Dox and subsequently sorted by FACS to create clonal cell lines, which were screened using immunofluorescence against Ki67.

### Single molecule RNA fluorescence in situ hybridization (smFISH)

Custom Stellaris RNA FISH probes labeled with Quasar dyes (i.e., Quasar®570 or Quasar®670) were designed against specific centromere RNAs and purchased from Biosearch Technologies (Petaluma, CA). To conduct single molecule FISH, cells grown on poly-L lysine coverslip in 12-well plates were washed with PBS and fixed with 4% paraformaldehyde in 1X PBS containing RVC (Ribonucleoside Vanadyl Complex) for 10 min at room temperature (RT). After washing cells twice with 1X PBS, cells were permeabilized in 70% ethanol for at least 20 min at 4°C. Cells were pre-incubated with 2X SSC; 10% deionized formamide for 5 min, and incubated with hybridization mix (0.1 μM RNA FISH, 10% deionized formamide, in Hybridization Buffer (Biosearch Technologies)) overnight at 37°C in the dark. Finally, cells were washed twice with 10% deionized formamide in 2X SSC for 30 min at 37°C and once with Wash B (Biosearch Technologies) for 5 min at RT. For experiments with immunofluorescence coupled to smFISH, HeLa cells grown on poly-L lysine coverslip in 12-well plates were washed with PBS and fixed with 4% paraformaldehyde in1X PBS containing RVC (Ribonucleoside Vanadyl Complex) for 10 min at RT. After washing cells with 1X PBS, cells were permeabilized for 5 min at RT with 0.1% Triton-X in PBS with RVC. After washing, primary and secondary antibody incubation was performed at RT for 1h in PBS and RVC. Antibody concentrations: DM1a (anti-tubulin; Sigma): 1:10000, ACA (anti-centromere antibodies – human auto-immune serum; Antibodies, Inc.): 1:1000, CENP-A (Abcam, ab13939) at 1:1000, Fibrillarin (Abcam, ab5921) at 1:300, and Ki67 (Abcam, ab155580) at 1:100. For smFISH, cells were fixed again for 10 min with 4% PFA in 1x PBS. Hybridization was performed as above. Coverslips were mounted on cells with Vectashield containing Hoechst.

For imaging, slides were imaged using a DeltaVision Core microscope (Applied Precision/GE Healthsciences) with a CoolSnap HQ2 CCD camera and 60x and 100x 1.40 NA Olympus U-PlanApo objective. smFISH foci were counted per nucleus from z-projected images using CellProfiler (Carpenter et al., 2006). Image files were processed using Deltavision software or Fiji (Schindelin et al., 2012).

**Supplemental Table 1. Probe sequences.** Also see Supplemental Table 2 for an analysis of matches to centromere reference sequences.

### ASAT Oligos

CAAAGAAGTTTCTGAGAATG

TGAGTTGAATGCACACATCA

AAAAGGAAGGTTCAACTCTG

TGTTTCAAAACTGCTCTATG

TGCAGATTCTACAAAAAGAG

AAGCGGTCCAAATATCCACT

CGTTTCCAACGAAGGCCTCA

TTTTTATATGAAGATATTCC

CTGAGAATGCTTCTGTCTAG

CACACATCACAAAGAAGTTT

TCAACTCTGTGAGTTGAATG

TGCTCTATGAAAAGGAAGGT

CAAAAAGAGTGTTTCAAAAC

ATATCCACTTGCAGATTCTA

AAGGCCTCAAAGCGGTCCAA

AGATATTCCCGTTTCCAACG

TCTGTCTAGTTTTTATATGA

### SF1 Oligos

TCTTTGTGGCCTTCGTTGGA

TCCAACGAAGGCCACAAAGA

GTTTCAAATCTGCTCTGTCT

AGACAGAGCAGATTTGAAAC

TTGTGGAATTTGCAAGTGGA

TCCACTTGCAAATTCCACAA

CTACGAAGGCCTCAAAGAGG

CCTCTTTGAGGCCTTCGTAG

AAATCTCCCCTTGCAAATTC

GAATTTGCAAGGGGAGATTT

GAAATCCCGTTTCCAACGAA

TTCGTTGGAAACGGGATTTC

ACCTTCCTTTAGACAGAGCG

CGCTCTGTCTAAAGGAAGGT

ATTTCCTTTTGTACCATTGG

CCAATGGTACAAAAGGAAAT

AAGTCTTTCCAAACTGCTCT

AGAGCAGTTTGGAAAGACTT

GTGTGCAAGTGGATATTTGG

CCAAATATCCACTTGCACAC

GTGCGTTCAACTCACAGAGT

ACTCTGTGAGTTGAACGCAC

TCACAAACAGAGGGTTTCCA

TGGAAACCCTCTGTTTGTGA

TTCCAACGAAGGCCTCAAAG

CTTTGAGGCCTTCGTTGGAA

GAGTGTTTCCAATCTACTCT

AGAGTAGATTGGAAACACTC

TGCAATGTGTGCGTTCAACT

AGTTGAACGCACACATTGCA

TGCTAGACAGAAGACTTCTC

GAGAAGTCTTCTGTCTAGCA

CCAATGGTAGAAAAGGAAAT

ATTTCCTTTTCTACCATTGG

GCAGTTTGGAAACACTCAGT

ACTGAGTGTTTCCAAACTGC

GTCAGCAACTGGATATTTGG

CCAAATATCCAGTTGCTGAC

ATTTGAGGCCTTCGTTGGAA

TTCCAACGAAGGCCTCAAAT

### SF3 Oligos

CAGCTGAAATTATCCCGTTT

AGTTCAACTCGGGGATTTGA

GGTCCAAATATCCACTTGTA

GATATACCCGTTTCGAACGA

TTCTGTCTAGATGGCATGTG

TGCAGTCATCACAGAGAAGC

CTGCGCTCTCAAAAGGAGAG

TGAAGATGATTCCGTTTCCA

TTCCTGAGAATGCATCTGTG

AGTTCAACTCTGGGAGTTGA

CATAGCTGCTCTTTCCAAAG

CAAACGTCCACTTGCAGATT

TTTTCCACCACAAACCACAA

GCAGTTTCTGAGAATGCTTC

TGTTCAACTCTGTGAGGTGA

GTTTCCAAACTGCTCTATCA

GCTTCTGTCTAGTTTTTCTA

GGTCCAAATATCCACTTGTA

TTATCCCGTTTCCAACGAAA

TCTGAGAATGCTTCCGTTTA

GATATACCCGTTTCGAACGA

TTTCTGAGAAGGCTTCTGTC

GGTTCAACTCTGTGAGTTGA

GAGAGTTTCAACACTGCTCT

ATATTCCCGTTTCCAAAGAC

CTCCAAATGTCCACTTGTAG

GTGACCATAATTCGTTTTCC

GTTTCTGACAATGCTTCTCT

GGAACCTTCAACTCTGTGAG

AATATCCACTTGCAGATTCC

GTTTAGCTTTCCTGTGAAGA

GTGTTTCAAAGCTTCTCTCT

ATATCTCCACTTGCAGATTC

AACTCAGTCGTCACCAAGAG

AAACTGCTCCATCCAAAGGA

TCCACTTGCACATTCTACAA

TTATCCCGTTTCCAACGAAA

ATGCTTCTGTTTAGTTCTGT

GAGCGGTCCAAATATCTACT

TTATCCCGTTTCCAACGAAA

ATGCTTCTGTTTAGTTCTGT

CATCACGAAGAGGGTTCTGA

GTTCAGCACTGTGAGTTGAA

ATATCCACTTGCAGTTTCTA

CGAAATGCTCAGAGAGGACC

AGTGCTTCTGTTTAGTTCTG

ACGGTACTCCTCAAAGAGTG

GCTCTGTGAGTTCAACTCAA

### ASAT Antisense Oligos

CATATAAAAACTAGACAGAA

GTTGGAAACGGGAATATCTT

TGGACCGCTTTGAGGCCTTC

AGAATCTGCAAGTGGATATT

TTTTGAAACACTCTTTTTGT

CCTTCCTTTTCATAGAGCAG

ATTCAACTCACAGAGTTGAA

AACTTCTTTGTGATGTGTGC

TAGACAGAAGCATTCTCAGA

GAATATCTTCATATAAAAAC

GAGGCCTTCGTTGGAAACGG

GTGGATATTTGGACCGCTTT

TCTTTTTGTAGAATCTGCAA

ATAGAGCAGTTTTGAAACAC

AGAGTTGAACCTTCCTTTTC

GATGTGTGCATTCAACTCAC

ATTCTCAGAAACTTCTTTGT

### X Oligos

TTGAATGCAGTCATCGCAGA

TCTGCGATGACTGCATTCAA

AACTGCTCCATCAGAAGGAT

ATCCTTCTGATGGAGCAGTT

TCCGAGAATGCTTCTGTTTA

TAAACAGAAGCATTCTCGGA

AATCTGCAAGTGGACGTTTG

CAAACGTCCACTTGCAGATT

GATTCCAATCTGCTCTATCA

TGATAGAGCAGATTGGAATC

TGGATATTTGGAGCTCTCTG

CAGAGAGCTCCAAATATCCA

GTGAACATATACCCGTTTCG

CGAAACGGGTATATGTTCAC

TGTTCAACTCTGGGAGTTGA

TCAACTCCCAGAGTTGAACA

AGCAGTTTCCAAACACACGT

ACGTGTGTTTGGAAACTGCT

### 7.2 Oligos

AGACTCTGCAAGTGGATATT

AATATCCACTTGCAGAGTCT

GCGCTTTGAAGCCTTCGTTG

CAACGAAGGCTTCAAAGCGC

ACTTCTTTGTGATGTGGGCA

TGCCCACATCACAAAGAAGT

CTTCAAAGCGCTCCAAATGT

ACATTTGGAGCGCTTTGAAG

GAAGATATTCCCTTTTCCAT

ATGGAAAAGGGAATATCTTC

TCTGTCTAGGTTTTAGGTGA

TCACCTAAAACCTAGACAGA

GTCGTTTCTGAGAATGCTTC

GAAGCATTCTCAGAAACGAC

ATTTGGAGCGCTTTGAAGAC

GTCTTCAAAGCGCTCCAAAT

TCAACTCACAGCGTTGAAAC

GTTTCAACGCTGTGAGTTGA

AGGTCGAAAAGGAAATATCT

AGATATTTCCTTTTCGACCT

## Supplemental Figure Legends

**Supplemental Figure 1. Centromere RNA levels vary across cell lines.** Related to Figure 1. Quantification indicating the variation of smFISH Foci across selected cell lines using probes designed against chromosome 3 (SF1). Error bars represent the mean and standard deviation of at least 100 cells.

**Supplemental Figure 2. Analysis of centromere RNAs following diverse perturbations.** Related to Figure 3. A) Treatment of HeLa cells with small molecule inhibitors against each polymerase reveals that alpha-satellite transcripts are mediated by RNA polymerase II. This experiment was conducted as in Figure 3A,B, but transcripts were detected with probe set designed against super-chromosomal family 1 sequences (SF1). B) Quantification of smFISH foci from (A) after treatment of HeLa cells with small molecule inhibitors against Cdk7, RNA Pol I and RNA Pol III. When compared to controls, the number of smFISH foci was substantially reduced after inhibition of RNA Pol II activator, Cdk7. Error bars represent the mean and standard deviation of at least 240 cells. C) Analysis of alpha-satellite smFISH transcripts (ASAT probe set) in cells depleted for diverse cell division components; proteins involved in centromere regulation (Sgo1 and BubR1), DNA replication (Mcm6, Gins1, Orc1, and Cdt1), sister chromatid cohesion (ESCO2, Scc1), chromosome condensation (Smc2, CAPG, CAPG2, TOP2A), and nucleosome remodeling (SSRP1) were targeted using Cas9. However, despite the diverse roles of these proteins in different aspects of centromere function, none of these inducible knockouts resulted in reduced levels of ASAT alpha-satellite transcripts as detected by smFISH analysis. Error bars represent the mean and standard deviation of at least 240 cells. D) The increase in alpha-satellite transcripts in cells depleted for CENP-C depends on RNA Polymerase II as THZ1 treatment resulted in a substantial reduction in smFISH foci in both control cells and CENP-C inducible knockout cells using SF1 probes. Scale bar, 25 µm.

**Supplemental Figure 3.** Related to Figure 4. A) Immunofluorescence of HeLa (top) and RPE-1 cell (bottom) showing the colocalization of centromeres with the nucleolus, as marked with antibodies against Fibrillarin and centromeres (ACA). B) Induction of Ki67 and Fibrillarin knockouts results in increased levels of alpha-satellite transcription, particularly for Ki-67, as tested by both ASAT and SF1 probe sets. The cell lines used for this experiment represent inducible knockout cells, in contrast to the stable Ki67 knockout analyzed in Fig. 4F,G. Error bars represent the mean and standard deviation of at least 240 cells. C) Validation of Ki67 stable knockout via immunofluorescence using antibodies against Ki67. Control cells (top) display clear Ki67 localization when compared to the clonal knockout cell line (bottom). Despite the persistently increased alpha-satellite transcript levels, there was no notable consequences to centromere protein levels (based on the localization of CENP-A). D) Graph showing quantification CENP-A intensity in control and Ki67 stable knockout cells. Each point represents the average of the centromeres within a single cell. N=15 cells/condition. Scale bars, 10 µm.

